# Adaptive coding of stimulus information in human frontoparietal cortex during visual classification

**DOI:** 10.1101/2021.11.22.469511

**Authors:** David Wisniewski, Carlos González-García, Silvia Formica, Alexandra Woolgar, Marcel Brass

## Abstract

The neural mechanisms of how frontal and parietal brain regions support flexible adaptation of behavior remain poorly understood. Here, we used functional magnetic resonance imaging (fMRI) and model-based representational similarity analysis (RSA) to investigate frontoparietal representations of stimulus information during visual classification under varying task demands. Based on prior research, we predicted that increasing perceptual task difficulty should lead to more categorical coding of stimulus information, and that exemplar-level stimulus coding would be restricted to posterior, sensory brain regions. Counter to our expectations, however, we found frontoparietal regions encoded exemplar-level stimulus information. Interestingly, the anterior intraparietal sulcus (aIPS) encoded stimuli equally well regardless of perceptual difficulty, and these representations were directly related to choice behavior (proportion of guessing). Overall, these findings reveal unexpected exemplar-level stimulus coding in frontoparietal cortex, and highlight the role of aIPS in supporting adaptive behavior.

## 1. Introduction

Regardless of the context we might find ourselves in, we can efficiently adjust our behavior to current demands ^1,2^, an ability that is supported by a set of frontal and parietal brain regions often called the multiple demand (MD) network ^3,4^. Past research using multivariate decoding ^5,6^ revealed that MD regions encode a wide range of task-related information, including stimuli, responses, and task-rules ^7^. Coding in MD regions is stronger for behaviorally relevant information that is explicitly attended ^8–10^, when performance is rewarded ^11^, or whenever tasks are particularly difficult ^12–14^. This has been taken as evidence for adaptive coding ^3^, i.e. the existence of multi-modal neurons rapidly and flexibly changing their coding properties to meet current demands. In turn these neurons are thought to bias coding in sensory and motor regions ^15,16^. However, other research demonstrated that, at least under some conditions, coding of task-related information in MD cortex does not adapt to changing demands ^17–19^, and instead the same representations are re-used regardless of the current context. Perceptual difficulty is a particularly interesting case here, since some researchers have suggested that MD cortex is unable to adapt to changes in the quality of perceptual input ^20^, while others have demonstrated adaptive coding under varying perceptual difficulties ^10,21^. Thus, it remains an open question whether adaptive coding is a general property of these regions, or whether frontoparietal cortex only adapts its representations under specific circumstances, and instead re-uses the same representations when necessary ^22–24^.

Previous research focused primarily on investigating *whether* coding adapts to different demands, for instance by testing whether information coding is stronger on hard than on easy trials ^12^, or stronger in freely chosen than in externally cued tasks ^18,25^. These studies have been optimized to detect the presence or absence of differences in coding strength across conditions, often using multivariate decoding methods ^6^, but make no explicit predictions about the representational formats used in each condition. Yet, changes to coding strength do not necessarily imply changes in coding format as well, and it remains difficult to draw strong conclusions about *how* representational formats change across conditions (e.g. more or less categorical coding). A complete explanation of how adaptive changes in neural coding are related to behavior requires both a description of changes to coding strength *and* format. We argue that understanding adaptive coding at the level of representational formats remains key to understanding the neural basis of goal-directed behavior.

Here, we used a visual classification task and assessed coding formats, to directly tackle this issue. In the past, visual classification was used successfully to study the representational format of task-related stimulus information, often in non-human primates ^26,27^. One of the key findings was that sensory and frontoparietal brain regions use different coding formats ^26^. Sensory regions preserved exemplar-level stimulus information, even if this was not behaviorally relevant ^28^. Frontoparietal brain regions only represented behaviorally relevant category information though, without representing exemplar-level information. This is usually interpreted as evidence for more abstract, behaviorally optimized stimulus coding in frontoparietal cortex ^27^. Here, we tested whether these results also obtained in population-level information coding in humans, and asked how these coding properties change when the perceptual difficulty of the task changed. Given the previous findings, we had three main hypotheses.

First, we expected that visual regions would preserve exemplar-level information on perceptually easy trials (which we call “equidistant coding” here since all stimuli used were equidistant in stimulus space). Increasing perceptual difficulty by adding noise to the stimuli was expected to weaken equidistant coding in visual regions (Hypothesis 1). Second, we expected to find stimulus and category coding in MD regions ^8,9,14^, but also expected a different effect of perceptual difficulty. Since coding of task-related information on MD regions is stronger on difficult trials ^10,12^, stimulus coding should be relatively weak on perceptually easy trials (Hypothesis 2), and stronger on perceptually difficult trials here as well. Third, and most importantly, we made explicit predictions about how representational formats might change across difficulty levels.

We expected representational formats to shift towards stronger categorical (“clustered”) coding on perceptually difficult trials (Hypothesis 3). Transforming exemplar-level to more categorical representations can be seen as a means to optimize these representations to support behavior in this classification task. From this perspective, on perceptually easy trials, MD regions might still contain some amount of exemplar-level information since there is little need for optimization and bottom-up stimulus information from sensory regions might be sufficient to correctly classify images. On perceptually difficult trials, the need to optimize representations is stronger, and behaviorally optimized categorical coding should thus be especially prevalent here. Such a shift in representational formats might be a key neural mechanism of how MD regions support adaptive behavior, especially under difficult conditions.

## 2. Results

In order to test these hypotheses, participants performed a simple classification task, determining whether morphed visual stimuli were either cats or dogs (morph level, from 100% cat to 100% dog). Perceptual difficulty was manipulated by adding random Gaussian noise to the images (noise level, clean vs noisy stimuli). Neural stimulus coding on clean and noisy trials in sensory and MD regions was assessed separately using a model-based representational similarity (RSA ^29,30^) approach.

### 2.1 Experimental task and behavioral results

Thirty-eight participants (29 female, 9 male) performed a visual classification task, which has been used successfully in the past ^26^, while undergoing fMRI. On each trial, a visual stimulus was presented on screen, and participants decided whether it was a cat or a dog (Figure 1A). Stimuli were morphed images, created from two different cat and dog “template” images, and ranging from 100% cat to 100% dog (morph level). This led some stimuli to be closer to the category boundary than others, effectively manipulating choice difficulty. Each image could be presented in one of two different perceptual difficult levels (clean = no noise, noisy = Gaussian noise added to stimulus, Fig 1B). While morph level was manipulated trial-by-trial, perceptual difficulty was manipulated block-wise. Each perceptual difficulty block started with the presentation of a verbal instruction on screen (“Easy/Hard block starting”). The amount of noise added was calibrated separately for each participant using a staircase procedure before the start of the experiment. The overall design was therefore 2(categories) × 2(noise levels) × 4(choice difficulty). Each trial consisted of the presentation of a single image on screen for 1.8s, followed by a variable inter-trial-interval (mean duration = 2.7s, range = 0.8 – 10.4s). Participants performed a total of six experimental runs of 128 trials each.

**Figure 1:**
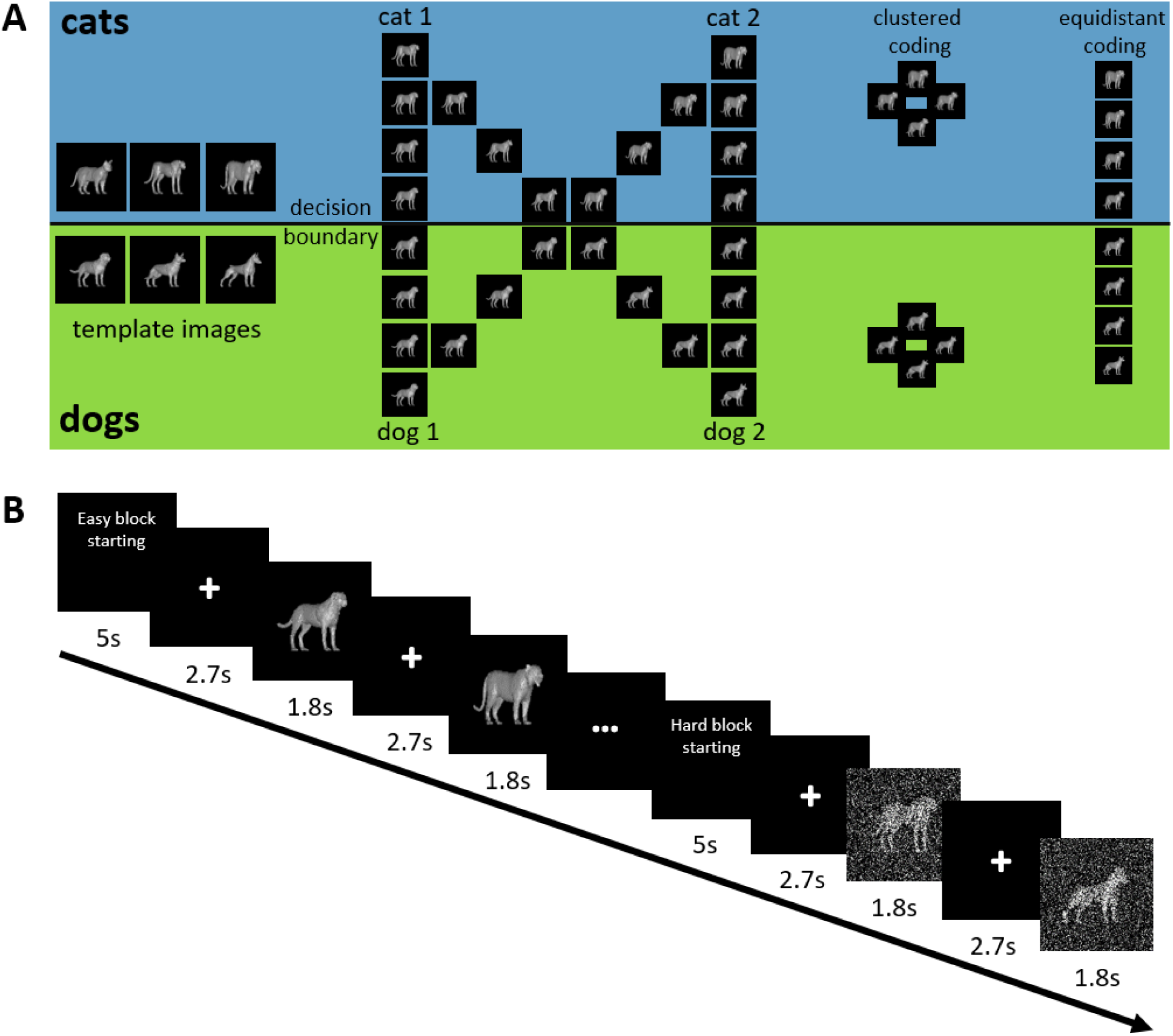
Experimental design. **A**. **Stimuli**. On the left, all template images used here are depicted. All stimuli above the decision boundary are cats, all below are dogs. Each participant was presented 2 out of 3 cat/dog templates (randomly selected for each participant). In the middle, example morphed stimuli are depicted for 2 dog and 2 cat template images. On the right, a schematic representation of clustered and equidistant coding is depicted. When stimulus representations are clustered, representational distances between exemplars of the same category (e.g. dog) are small, i.e. representations are highly similar. Distances between categories are large. When stimulus representations are equidistant, distances from the perceptual space (middle) are preserved in the neural space, including differences between exemplars within the same category. **B**. **Trial timing**. Each block started with an instruction screen. Then, stimuli were presented for 1.8s, interleaved with a variable inter-trial-interval (mean duration 2.7s, range 0.8 – 10.4s).

We first characterized behavioral performance by collapsing data across both categories, and then computing a 2 (noise level) × 4 (choice difficulty) Bayesian ANOVA on the error rates (Figure 2A, left panel). We found performance to range from 2.33% errors to 44.25% errors, and found very strong evidence for main effects of both noise level and choice difficulty, BF10s > 150, with noisy trials and high choice difficulty trials having higher error rates. The effect of noise level decreased with increasing choice difficulty (interaction effect, BF10 > 150), likely reflecting a floor effect in performance, with classification on stimuli very close to the decision boundary being so difficult that adding noise had a negligible effect on performance. We found similar results for reaction times (Figure 2A, right panel), with very strong evidence for both main effects, BF10 > 150, and strong evidence for an interaction effect, BF10 = 80.51. Only correct trials were used in RT analyses.

**Figure 2.**
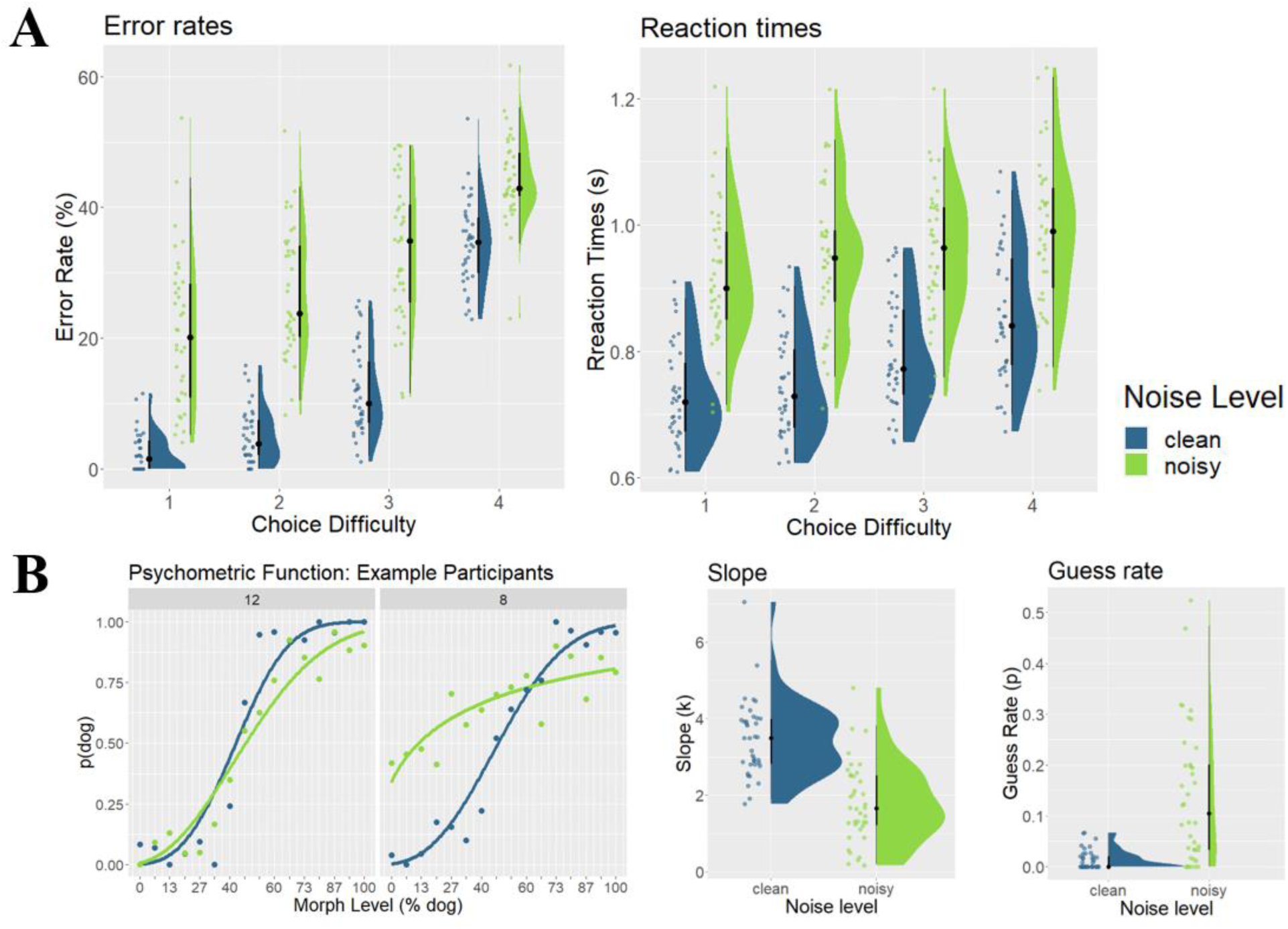
Behavioral Results. **A**. **Error rates and reaction times**. Plots show error rates (left) and reaction times (right) as a function of choice difficulty (1 = easy / far away from decision boundary, 4 = hard / close to decision boundary). Raincloud plots include boxplots centered around the median (black lines), probability density estimates (right half), and raw data (left half), jittered for illustration purposes. **B. Psychometric functions**. Example choice data from two participants (#12, #8, left). Each dot represents the probability of choosing ‘dog’ (p(dog)), as a function of morph level (expressed in % dog included in the morphed image). Lines represent fitted psychometric functions. The performance of one participant (#8) was heavily affected by noise level, while it was not for the other participant (#12). Estimated slope (k) and guess rate parameters are shown on the right, as a function of noise level (clean, noisy). blue = clean trials, green= noisy trials.

Next, we assessed performance by fitting psychometric functions to the choice data of each participant (Figure 2B). We first tested whether the slope of the function was higher on clean than on noisy trials, which would indicate a sharper category separation. We found very strong evidence for this effect, BF10 > 150 (mean slope_clean_ = 3.48, mean slope_noisy_ = 1.81). We then tested whether choices were biased towards either cats or dogs by analyzing the threshold parameter of the psychometric function, and found no evidence for biased choices in either clean or noisy trials (Supplementary Analysis 1). Lastly, we investigated whether the guess rate (proportion of trials in which participants guessed) was higher on noisy than on clean trials. We found very strong evidence for this effect, BF10 > 150 (mean guess_clean_ = 0.01, mean guess_noisy_ = 0.13), showing that overall, adding noise led to substantially more guessing, and a weaker separation of both categories.

### 2.2 Model based representational similarity analysis reveals stimulus coding in perceptual and frontoparietal brain regions

In order to investigate the adaptive coding of stimulus information in frontoparietal cortex, we employed a model-based representational similarity approach (RSA, ^29,30^, Figure 3, middle and right panels). For each region-of-interest (ROI, Figure 4), we first extracted parameter estimates for each of the 16 different conditions in this task (2 categories × 2 noise levels × 4 choice difficulty levels). Then, we computed pair-wise cross-validated Euclidean distances, which were represented in a 16×16 representational distance matrix (RDM, Figure 3, left panel).

**Figure 3:**
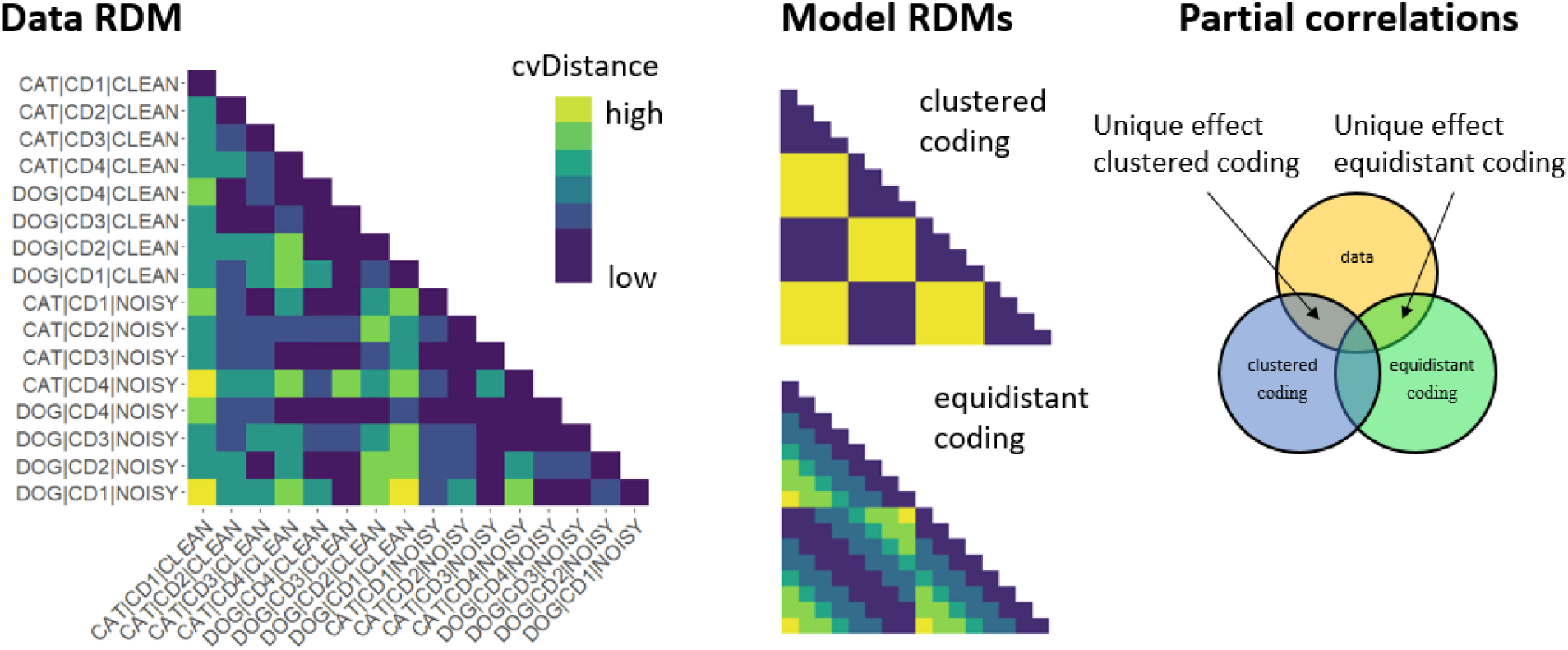
Model-based RSA. On the left, an example data representational distance matrix (RDM) is depicted, from the anterior intraparietal sulcus (IP2). Each of the 16 conditions (2 categories: cat, dog; 4 choice difficulty levels: cd1, cd2, cd3, cd4; 2 noise levels: clean, noisy) is represented in one row / column. Each cell represents the pairwise cross-validated Euclidean distance (cvDistance) between beta vectors from the corresponding conditions. Dark colors indicate low distances, while bright colors indicate high distances. In the middle, two stimulus coding models are represented in the same RDM format as the data. On the right, a schematic representation of our partial correlation analysis demonstrates how unique shared variance between each model and the data RDM was computed.

**Figure 4:**
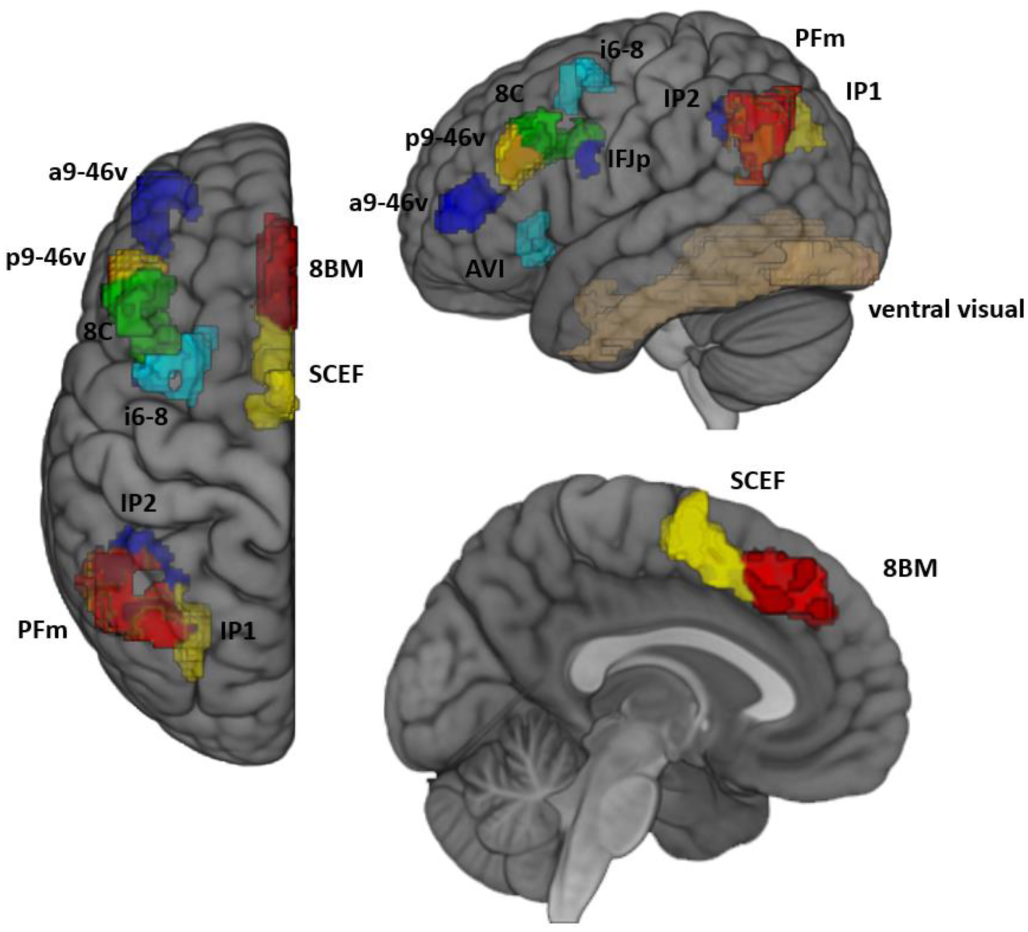
ROIs. ROIs were derived from the HCP atlas ^31^ and included the multiple demand regions identified in ^32^. The ventral visual cortex ROI was defined using the Harvard-Oxford atlas.

We defined two alternative theoretical models of how stimulus information could be encoded in the brain: equidistant and clustered coding. First, neural representations could preserve distances between individual exemplars in stimulus space (isomorphic coding, which we call ‘equidistant’ coding here since stimuli were equidistant in stimulus space). Equidistant coding (Figure 1A) fully preserves the stimulus space and differentiates between individual exemplars within a category, even though this information is not relevant to the categorization task. We generated a model RDM from this model, and then computed canonical, zero-order correlations between the model and data RDMs, separately for each ROI and participant. Positive correlations indicate that the equidistant coding model explained data variance in a given ROI. Second, neural representations could be more categorical and cluster stimulus representations of individual exemplars within the same category (clustered coding), making neural distances within categories small and between categories large (Figure 1A). Clustered coding is thus optimized for behavioral performance, since it collapses across task-irrelevant stimulus features (exemplars within categories) and only contains task-relevant category information. We again generated a model RDM here (Figure 3), and then correlated model and data RDMs to quantify whether clustered coding explains a part of the data variance. Note that both of these models would be indistinguishable using the design and multivariate decoding methods from past research, since these could, in principle, successfully differentiate cats and dogs regardless of whether representations were equidistant or clustered.

As expected, results indicated that on clean trials, ventral visual cortex showed strong evidence for both equidistant (r = 0.14, BF10 > 150) and clustered coding (r = 0.08, BF10 = 80.00, Figure 5, see Table 1 for all results). In MD cortex, we found evidence for equidistant coding in posterior dlPFC (8C, r = 0.05, BF10 = 4.89), and IPS (IP1, r = 0.06, BF10 = 7.49, IP2, r = 0.09, BF10 > 150). There was anecdotal evidence for equidistant coding in the anterior dlPFC (p946v, r = 0.04, BF10 = 2.81) and the angular gyrus (PFm, r = 0.05, BF10 = 2.85). No MD ROI showed evidence for clustered coding (all BF10s < 0.58). Thus, we were able to detect stimulus information in visual, parietal, and lateral prefrontal brain regions on clean trials. On noisy trials, we found strong evidence for equidistant coding in the posterior IPS (IP1, r = 0.06, BF10 = 28.14) and moderate evidence for clustered coding in the same region (r = 0.05, BF10 = 8.25). The anterior IPS showed anecdotal evidence for equidistant coding on noisy trials (IP2, r = 0.06, BF10 = 2.85). We also found evidence for equidistant coding in dmPFC (8BM, r = 0.06, BF10 = 5.90). Importantly, it remains difficult to attribute results to a specific coding model here, since both candidate models were positively correlated and zero-order correlations do not explicitly control for their covariance. These results demonstrate however that, in principle, we are able to detect both coding formats on both clean and noisy trials.

**Figure 5.**
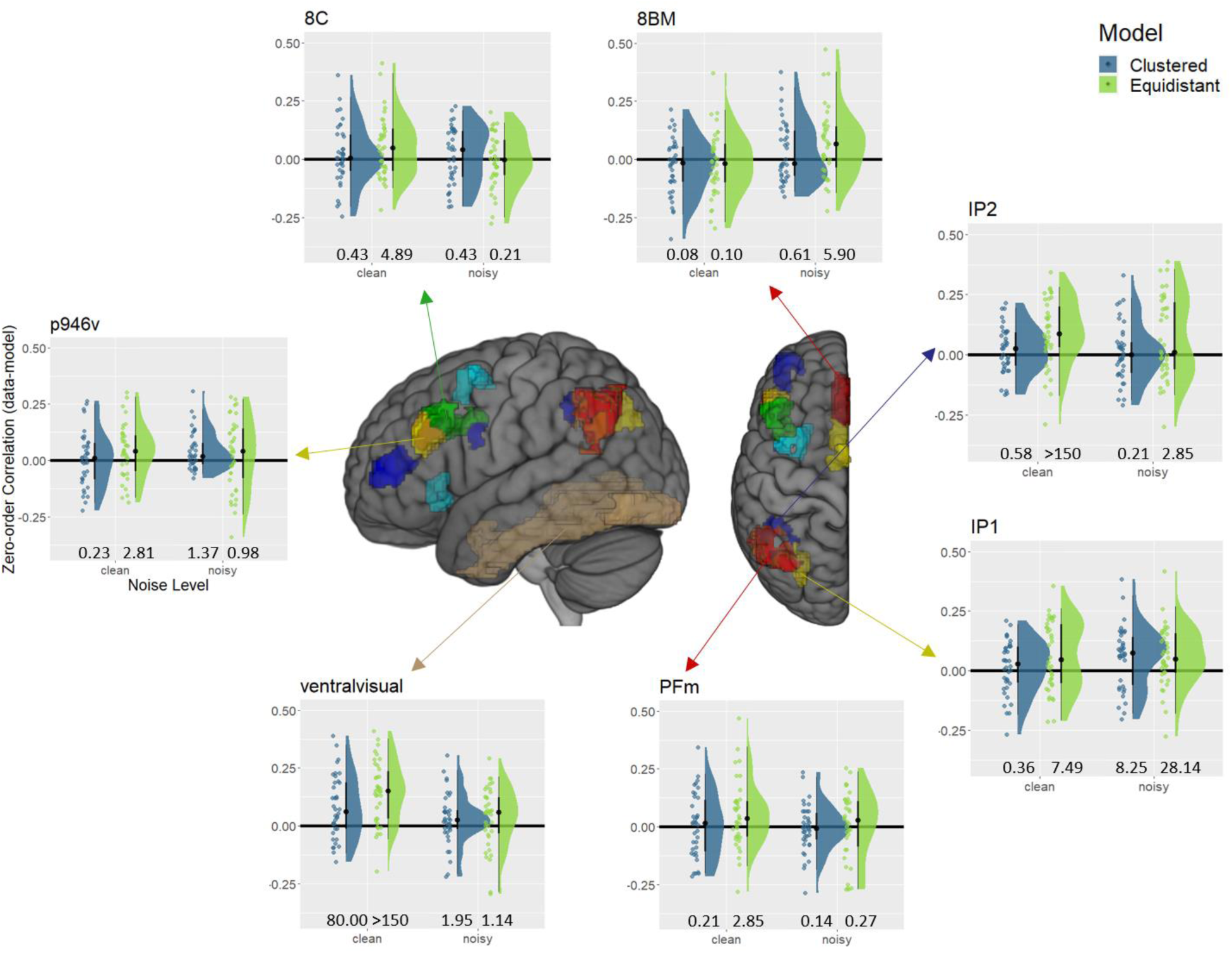
Model-based RSA results – zero-order correlations. Raincloud plots ^33^ depict the canonical / zero-order correlation of both models (clustered = blue, equidistant = green) with the data RDM of each ROI. Raincloud plots include boxplots centered around the median (black lines), probability density estimates (right half), and raw data (left half), jittered for illustration purposes. Numbers at the bottom of the plots indicate Bayes factors of a t-test against zero. We only depict results from ROIs that showed evidence for an effect in at least one of the four conditions depicted.

**Table 1:**
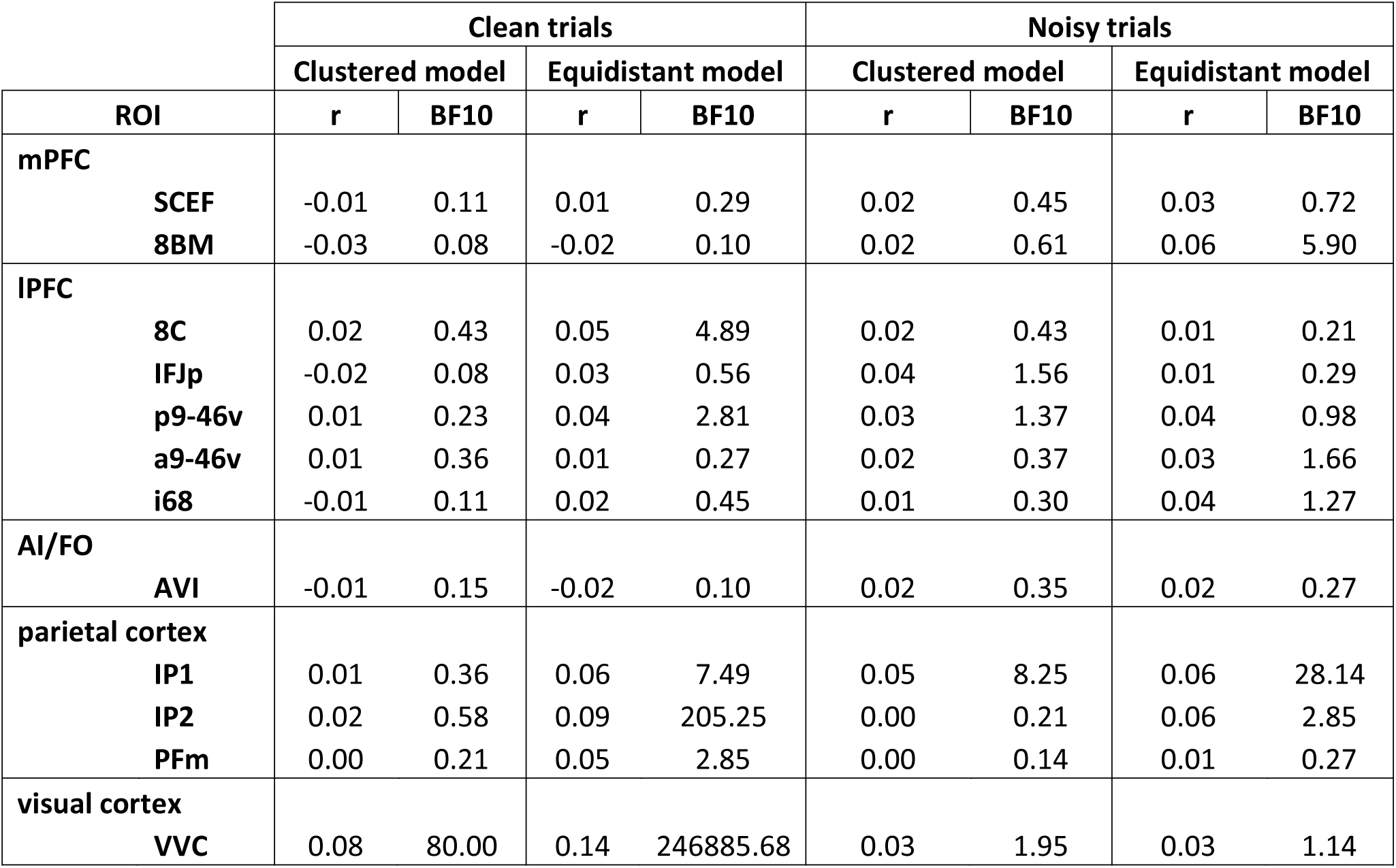
Zero-order correlations. Canonical / zero-order Spearman correlations (r) between the models and the data RDM, including the Bayes factor (BF10) of the corresponding t-test against zero, separately for clean and noisy trials. mPFC = medial prefrontal cortext (SCEF, 8BM), lPFC = lateral prefrontal cortex (8C, IFJp, p9-46v, a9-46v, i68), AI/FO = anterior insula / frontal operculum (AVI), VVC = ventral visual cortex

### 2.3 Visual cortex retains exemplar-level stimulus information under perceptually easy conditions

In order to determine whether the equidistant or clustered models explained a unique part of the data variance, we repeated the model-based RSA outlined above, only now using partial instead of canonical correlations. Crucially, partial correlations allowed us to remove any variance that the data RDM shared with both models (Figure 3), and only investigate the unique shared variance between e.g. the equidistant model and the data RDM, while controlling for variance shared with the clustered coding model.

In ventral visual cortex, we expected the equidistant coding model to explain a unique part of the data variance on clean trials (Hypothesis 1). Indeed, we found very strong evidence for that hypothesis (r = 0.12, BF10 > 150, Figure 6, see Table 2 for full results). Interestingly, for noisy trials, we found moderate evidence against a unique contribution of equidistant coding (r = 0.02, BF10 = 0.33). Directly comparing both results using a paired t-test yielded moderate evidence for a difference (BF10 = 4.97), suggesting that the equidistant coding model better explained data on clean than on noisy trials. These results are largely in line with our predictions. Ventral visual cortex indeed encoded the full stimulus space using an equidistant format on clean trials. This signal became weaker on noisy trials, likely reflecting the fact that stimulus input was degraded.

**Figure 6.**
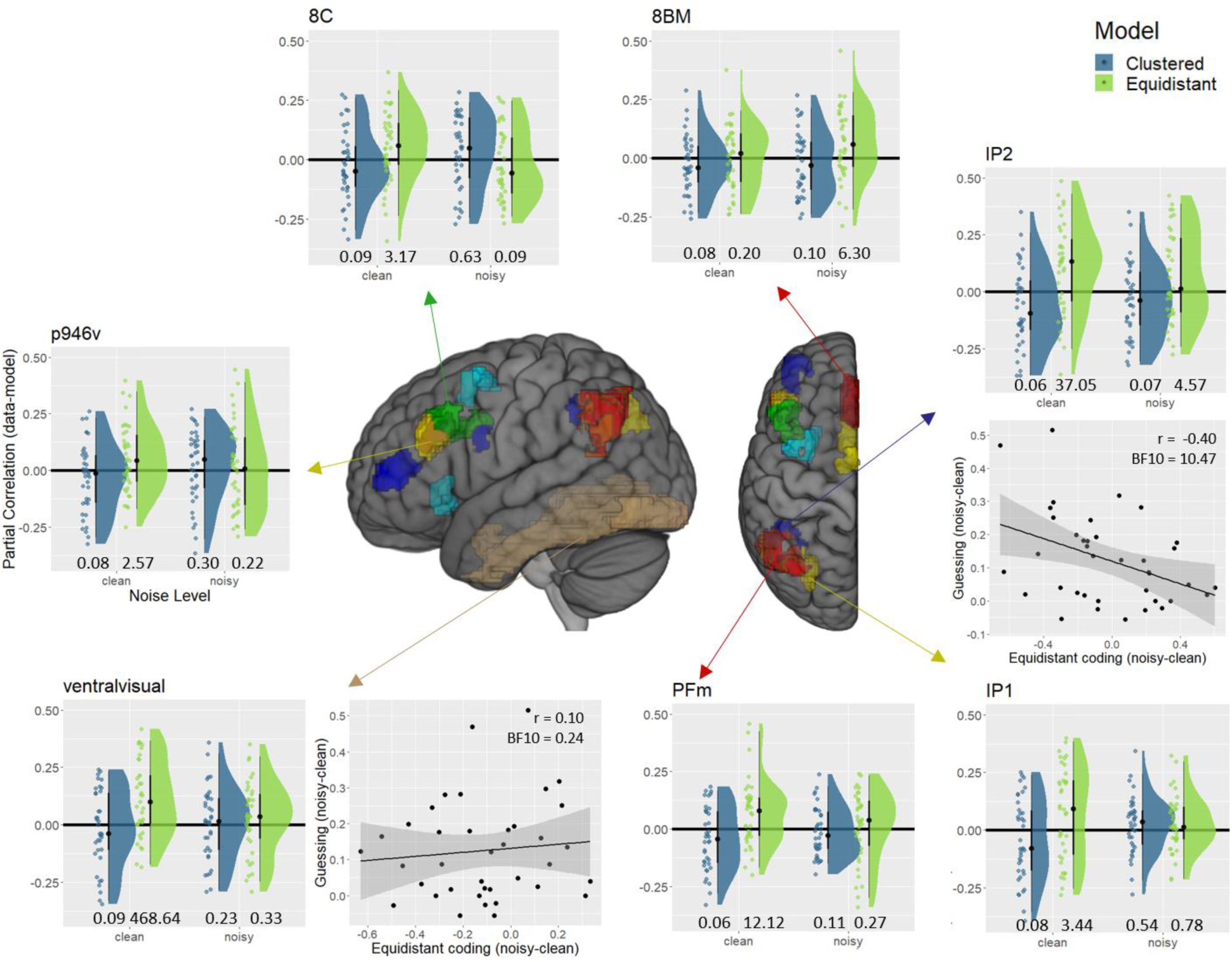
Model-based RSA results – partial correlations. Raincloud plots ^33^ depict the partial correlations of both models (clustered = blue, equidistant = green) with the data RDM of each ROI, controlling for the other model, respectively. Raincloud plots include boxplots centered around the median (black lines), probability density estimates (right half), and raw data (left half), jittered for illustration purposes. Numbers at the bottom of the plots indicate Bayes factors of a t-test against zero. We only depict results from ROIs that showed evidence for an effect in the manipulation check (Figure 5). For ventral visual cortex and IP2, we also depict brain behavior correlations. The x-axis shows the difference in equidistant coding between noisy and clean trials. The y-axis shows the difference in guessing, as estimated using psychometric functions, between noisy and clean trials. In IP2, the more equidistant coding collapsed in noisy trials, the more participants guessed. This was not the case in ventral visual cortex.

**Table 2:** Partial correlations. Partial Spearman correlations (r) between the models and the data RDM, including the Bayes factor (BF10) of the corresponding t-test against zero, separately for clean and noisy trials. mPFC = medial prefrontal cortext (SCEF, 8BM), lPFC = lateral prefrontal cortex (8C, IFJp, p9-46v, a9-46v, i68), AI/FO = anterior insula / frontal operculum (AVI), VVC = ventral visual cortex

|  |  | Clean trials |  |  |  | Noisy trials |  |  |  |
| --- | --- | --- | --- | --- | --- | --- | --- | --- | --- |
|  |  | Clustered model |  | Equidistant model |  | Clustered model |  | Equidistant model |  |
| ROI |  | r | BF10 | r | BF10 | r | BF10 | r | BF10 |
| mPFC | SCEF | -0.03 | 0.08 | 0.03 | 0.58 | -0.01 | 0.14 | 0.03 | 0.46 |
|  | 8BM | -0.03 | 0.08 | 0.00 | 0.20 | -0.02 | 0.10 | 0.06 | 6.30 |
| IPFC | 8C | -0.02 | 0.09 | 0.06 | 3.17 | 0.03 | 0.63 | -0.03 | 0.09 |
|  | IFJp | -0.06 | 0.06 | 0.06 | 2.82 | 0.05 | 2.38 | -0.03 | 0.10 |
|  | p9-46v | -0.03 | 0.08 | 0.05 | 2.57 | 0.02 | 0.30 | 0.01 | 0.22 |
|  | a9-46v | 0.01 | 0.22 | 0.00 | 0.19 | 0.00 | 0.20 | 0.02 | 0.39 |
|  | i68 | -0.04 | 0.07 | 0.05 | 1.36 | -0.03 | 0.08 | 0.06 | 1.78 |
| AI/FO | AVI | 0.01 | 0.27 | -0.02 | 0.10 | 0.01 | 0.24 | 0.01 | 0.24 |
| parietal cortex | IP1 | -0.04 | 0.08 | 0.07 | 3.44 | 0.02 | 0.54 | 0.03 | 0.78 |
|  | IP2 | -0.07 | 0.06 | 0.11 | 37.05 | -0.05 | 0.07 | 0.08 | 4.57 |
|  | PFm | -0.04 | 0.06 | 0.07 | 12.12 | -0.01 | 0.11 | 0.01 | 0.27 |
| visual cortex | VVC | -0.03 | 0.09 | 0.12 | 468.64 | 0.01 | 0.23 | 0.02 | 0.33 |

As a control, we repeated the analysis assessing evidence for a unique contribution of the clustered coding model. We found moderate evidence against coding model explaining a unique part of the data variance on clean trials (r = -0.03, BF10 = 0.09), and the same was true for noisy trials as well (r = 0.01 BF10 = 0.23). The paired t-test yielded moderate evidence against a difference between these values (BF10 = 0.29).

### 2.4 Frontoparietal cortex shows weak stimulus coding under perceptually easy conditions

We then focused our analyses on MD regions, expecting to see relatively weak stimulus coding on perceptually easy trials (Hypothesis 2). To test this hypothesis, we used the same partial correlation approach outlined in the analysis above. We restricted the number of ROIs to those that showed some evidence for an effect in the zero-order correlation analysis, since it can be difficult to interpret partial correlations in the absence of zero-order correlations. This included parietal cortex (IP1, IP2, PFm), dlPFC (8C, p946v), and dmPFC (8BM). We found evidence for a unique contribution of equidistant coding in parietal cortex (IP1, r = 0.07, BF10 = 3.44, IP2, r = 0.11, BF10 = 37.05, PFm, r = 0.07, BF10 = 12.12, Figure 6), and dlPFC (8C, r = 0.06, BF10 = 3.17, p946v, r = 0.05, BF10 = 2.57), but not in dmPFC (8BM, r = 0.00, BF10 = 0.20). As a control analysis, we also tested for evidence for a unique contribution of clustered coding. For all MD ROIs, we found evidence against clustered coding (all rs < 0.01, all BF10s < 0.27), and this model thus failed to explain unique variance on perceptually easy trials in all ROIs assessed here. Overall, parietal cortex seemed to robustly represent the full stimulus space on clean trials, with effects in dlPFC being somewhat weaker. These findings were largely in line with our predictions, MD stimulus coding on clean trials was either absent, too weak to be detected reliably, or present only in an equidistant format that might indicate bottom-up stimulus information being passed on from ventral visual cortex.

### 2.5 Frontoparietal cortex encodes exemplar-level stimulus information under perceptually difficult conditions

The key prediction in this study was that we would observe adaptive stimulus coding in frontoparietal cortex. Specifically, we hypothesized that clustered coding would be strengthened on noisy trials as a means to compensate for the increased task difficulty (Hypothesis 3). In contrast to equidistant coding, clustered coding removes unnecessary exemplar-level information from neural representation, and instead focuses solely on the behaviorally relevant stimulus dimension (i.e. category). Counter to our expectations, we only found anecdotal evidence for a unique contribution of clustered coding on noisy trials in IFJp (r = 0.05, BF10 = 2.38). For all other ROIs, we found evidence against such an effect (all rs < 0.04, all BF10s < 0.63, see Table 2 for full results). Paired t-tests revealed that only IFJp showed evidence for stronger clustered coding on noisy as compared to clean trials (BF10 = 9.50, all other BF10s < 1.05). Thus, of all MD ROIs, only IFJp shows an effect that was in the expected direction, but this finding remains difficulty to interpret since we found no strong evidence for a corresponding zero-order correlation with the clustered model in IFJp (Table 1).

Unexpectedly, we found evidence for a unique contribution of equidistant coding on noisy trials in parietal cortex (IP2, r = 0.08, BF10 = 4.57), as we had seen in this region on clean trials above (IP2, r = 0.11, BF10 = 37.05). We also found evidence for equidistant coding in dmPFC on noisy trials (8BM, r = 0.06, BF10 = 6.30), which has shown evidence against equidistant coding on clean trials (8BM, r = 0.00, BF10 = 0.20). Thus the numerical trend was for equidistant coding to decrease on noisy trials in IP2, and to increase in 8BM. However, paired t-tests revealed moderate evidence against differences across noise levels in IP2 (BF10 = 0.32) and anecdotal evidence against a difference in 8BM (BF10 = 0.80), indicating that these differences were not statistically reliable.

One possible explanation for the relative lack of clustered stimulus coding in MD regions on noisy trials might be that the signal-to-noise-ratio was too low to reliably detect stimulus information. We believe this explanation to be unlikely though, as we were able to detect equidistant stimulus coding in parietal and medial prefrontal cortex on noisy trials. It has been shown previously that detecting task-related information in prefrontal cortex is especially difficult ^34^, and the fact that we were able to detect stimulus information in prefrontal cortex on noisy trials shows that, in principle, experimental power is high enough to detect such information even in small regions where signals are often weak.

Another possible explanation might be that our analysis approach was less sensitive to detect clustered coding, since this model had an overall lower variance / complexity than the equidistant model and might therefore be harder to detect in (partial) correlation analyses. To directly assess this possibility, we performed a number of simulation analyses. Using simulated data, we demonstrate that if anything, our analysis approach was *more* sensitive to detect clustered coding, making our results even more surprising (Supplementary Analysis 2).

### 2.6 Equidistant coding in anterior IPS is related to choice behavior

In additional exploratory analyses, we tested whether equidistant stimulus coding in anterior IPS (IP2), dmPFC (8BM) or ventral visual cortex was related to choice behavior. For this purpose, we correlated the partial correlations between the equidistant model and data RDMs with the parameters extracted from the psychometric function (slope, guess rate).

First, we reasoned that a sharper category boundary, as operationalized through the slope of the psychometric function, would be positively related to stimulus coding. We only investigated the anterior IPS here, since we found evidence against stimulus coding on clean trials in the dmPFC. In anterior IPS, we found evidence against a relationship between slope and stimulus coding, r = -.012, BF10s = 0.23. One potential explanation for this finding might be that using partial correlations, we remove at least some of the variance related to object categories, making it unlikely to find a relation to sharper category distinctions in behavior. Repeating the same analysis using zero-order correlations yielded similar results however, r = -0.04, BF10 = 0.30, making this explanation unlikely.

Second, we reasoned that weak stimulus signals would lead to fewer decisions that are based on stimulus evidence, and thus lead to more guessing. Correlating equidistant coding (after partialling out clustered coding effects) with guess rate on clean trials yielded no evidence for a relation between these variables (ventral visual, r = -0.01, BF10 = 0.37, IP2, r = -0.03, BF10 = 0.42, 8BM, r = -0.23, BF10 = 1.46). Interestingly, we found a relationship between equidistant coding and guessing specifically on noisy trials in IP2 (r = -0.39, BF10 = 9.81), but not in ventral visual cortex (r = 0.13, BF10 = 0.22) or 8BM (r = 0.06, BF10 = 0.28). Weaker equidistant coding on noisy trials in IP2 was associated with more guessing, and this effect was specific to noisy trials only (95% CI_r(clean)_ = [-0.63,-0.09], r_noisy_ = -0.03).

In order to test more directly whether an increase in guessing from clean to noisy trials was related to a decrease in equidistant stimulus coding, we computed difference scores between noise levels for both measures for each participant (r(clean) – r(noisy), guess(clean) – guess(noisy)). Correlating these difference scores yielded evidence for a negative correlation in IP2 (r = -0.40, BF10 = 10.47, Figure 6), indicating that participants with a weaker decrease in equidistant stimulus coding on noisy trials also showed a smaller drop in performance on these trials. We found evidence against such a relationship in ventral visual cortex (r = 0.10, BF10 = 0.24), and 8BM (r = -0.01, BF10 = 0.37).

## 3. Discussion

In summary, we found evidence for stimulus coding in a visual classification task in both parietal cortex (IP1, IP2, PFm), dlPFC (8C, p9-46v) and dmPFC (8BM). We expected to see evidence for adaptive coding of stimulus information in these regions, with stronger and more categorical coding under perceptually difficult, as compared to easy conditions. While we found weak evidence for stronger coding in some brain regions (dmPFC), we found no evidence for the proposed shift toward more categorical coding. On the contrary, our results suggest that frontoparietal cortex encoded the full stimulus space (equidistant coding), preserving behaviorally irrelevant exemplar-level stimulus information, under both perceptually easy and difficult conditions. The anterior IPS (IP2) showed a particularly interesting pattern of results, as equidistant stimulus coding here was equally strong under perceptually easy and difficult conditions, in contrast to e.g. ventral visual cortex, which showed a marked decrease in stimulus coding. Coding in the anterior IPS was also related to choice behavior, with stronger equidistant stimulus coding being related to less guessing.

Past research on perceptual decision-making and visual classification demonstrated that both visual and prefrontal brain regions encode stimulus information ^8,9,26^. Crucially, while ventral visual cortex represents the full stimulus space, i.e. information about both categories and individual exemplars ^28,35^, higher-level brain regions such as the prefrontal cortex have been reported to carry little information about individual exemplars in both humans ^37^ and non-human primates ^26,36^. These findings demonstrate a tuning of “perceptual” representations towards the task goal, successful categorization. Such abstract, “behavioral” representations only maintain task-relevant information (categories) and remove task-irrelevant information (exemplars). Our results in ventral visual cortex are largely in line with this research. We found stimuli to be encoded in an equidistant format that preserves information about the full stimulus space, and coding was stronger on perceptually easy than difficult trials. This indicates that ventral visual cortex shows “perceptual” stimulus coding that is weakened when stimulus input is degraded.

In MD cortex, we expected to see evidence for behavioral stimulus coding, i.e. categorical stimulus representations, especially under perceptually difficult conditions. In addition to previous findings, this hypothesis was informed by recent theories on adaptive coding ^24,38^. The transformation of perceptual into behavioral stimulus representations makes coding more categorical ^27^, and this essentially reflects a dimensionality reduction in neural representations. High-dimensional perceptual representations that carry information about the whole stimulus space are transformed into low-dimensional, behavioral representations that merely carry information about the task-relevant stimulus dimension. Recent theories suggest that high-dimensional representations are more easily separable, but also more susceptible to noise ^24,38^. Low-dimensional representations are more robust to noise, suggesting they would be especially useful on noisy, perceptually difficult trials, where stimulus information is degraded.

Our results in MD cortex were unexpected, and not in line with these predictions. We observed equidistant stimulus coding on perceptually easy trials in parts of the MD network, (parietal cortex and dlPFC, but not dmPFC and anterior insula / frontal operculum). On perceptually difficult trials, we found no strong evidence for clustered stimulus coding. There was anecdotal evidence for such an effect in the inferior frontal junction (IFJp), but this was difficult to interpret since it was based on partial correlation results in the absence of a corresponding zero-order correlation. Surprisingly, we did find evidence for *equidistant* stimulus coding that preserved exemplar-level information in the aIPS and dmPFC. There are two notable aspects to this finding. First, we were unable to find stronger coding on difficult as compared to easy trials, which has been observed before in MD cortex for stimulus position ^12^, stimulus shape ^10^ and task rules ^13^. On the contrary, information coding in the MD system tended to decrease for noisy trials, except in aIPS in which coding was equally strong on both easy and hard trials. Second, instead of showing the predicted shift from equidistant to clustered stimulus coding, aIPS instead encoded stimuli in an equidistant format on both perceptually easy and difficult trials.

One potential explanation for this might be the overall difficulty of the classification task. Error rates were over 30% for stimuli close to the decision boundary, and it might be that stimulus and category signals were very weak as a result. Although we expected precisely this fact to drive clustered stimulus representations, we cannot rule out that these representations were too weak to detect in some cases. However, our simulations (Supplementary Analysis 2) demonstrated that, if anything, our analysis was more sensitive to finding clustered coding, and less sensitive to finding equidistant coding. Thus, having found evidence for equidistant coding indicates that we could, in principle, have found evidence for clustered coding as well.

Another, more interesting, explanation for this finding is that it might be driven by the type of experimental manipulation we used in this study. By definition, MD regions are more strongly engaged in difficult, as compared to easy tasks in multiple domains ^3,4,32^. Recently, it has been suggested, however, that some types of difficulty manipulation might engage MD regions more than others ^20^. Specifically, difficulty manipulations that limit or degrade incoming stimulus information (e.g. increased noise) might not recruit MD regions as much as manipulations that make stimulus processing harder (e.g. mental rotation ^39^). This would make sense if MD regions recruit additional attentional resources to compensate for increased processing demands ^40^, but are not involved in refining stimulus representations themselves. From this perspective, degrading stimulus information might be a difficulty increase that MD regions cannot compensate for, and are not strongly involved as a result. Obviously, this interpretation is speculative at the moment, and does not immediately account for all the existing literature ^10^, but could account for the findings reported here. Future research using different, more processing-based difficulty manipulations will be needed to directly compare to these results.

Of all regions assessed in this study, the aIPS appeared to have a special role in compensating for increased perceptual difficulty. aIPS is involved in the processing of stimulus features ^41^, but also more broadly implied in regulating top-down attention ^42^. Previous work in non-human primates further implies that stimulus representations are not as categorical as in e.g. dlPFC ^27^, which might help explain why we saw equidistant coding here. Yet, it was the only brain region that consistently encoded stimulus information on both clean and noisy trials. This is in contrast to the ventral visual cortex, which only encoded stimuli on clean, but not on noisy trials. Thus, aIPS contained stimulus information under perceptually difficult conditions that was undetectable in perceptual brain regions, suggesting a role in maintaining performance under challenging conditions. It would have been reasonable to predict a similar pattern of results in e.g. dlPFC, even though effects here are often weaker ^34^, but we only found this results in aIPS. One potential interpretation might be that aIPS maintains attention to relevant stimulus features under varying difficulties, and thus enables meaningful, evidence-based decisions even under difficult conditions. This would also help explain why aIPS represents stimuli in an equidistant format. Since stimulus features differ between different exemplars within the same category, it would be sub-optimal to discard exemplar-level information and only encode category information instead. Maintaining exemplar-level information instead allows the aIPS to direct attention towards any relevant feature, under perceptually easy or difficult conditions. This interpretation is further supported by finding a correlation between stimulus coding in aIPS and choice behavior, specifically on perceptually difficult trials. Clearly, more research will be needed to replicate this exploratory finding, yet it suggests that aIPS is important for adapting to difficult conditions in a perceptual decision-making task.

In conclusion, we observed stimulus coding in a visual classification task in visual and frontoparietal brain regions. Unexpectedly, stimulus coding in frontoparietal regions retained exemplar-level stimulus information, despite it being task-irrelevant. Counter to our predictions, increasing perceptual difficulty did not make stimulus coding more categorical. We found some evidence for adaptive coding nonetheless, with aIPS encoding stimuli equally well under perceptually easy and difficult conditions, despite weaker input from visual cortex, in a manner that predicted behavior. These results reveal how specific representational formats do and do not change under varying task demands, and thus provide novel evidence of *how* adaptive neural coding in frontoparietal cortex supports adaptive behavior.

## 4. Methods

### 4.1 Participants

Forty-nine volunteers (38 female, 11 male, mean age: 24.1 years, range: 18-36 years) with normal or corrected-to-normal vision participated in the study. We obtained written informed consent from each participant prior to participation, and the Ethics Committee of the Ghent University Hospital approved this experiment (project identifier BC-07446). Each volunteer received 43€ for their participation. We first calculated the average error rate for each participant in the easiest possible condition in this experiment (clean template images, see below for more details). Five participants had excessive error rates (above 1.5*IQR of the group mean,): 10.4%, 10.4%, 12.5%, 16.7%, and 22.9% respectively, group average = 3.1%. These participants were excluded from the sample. Four participants showed excessive head movement during scanning (> 5mm), and were also removed. Two additional subjects were removed due to technical difficulties. The final sample consisted of thirty-eight participants (29 female, 9 male, mean age = 24.4 years, range: 19-36 years).

### 4.2 Task: Stimuli and Design

Stimuli consisted of gradually morphed, greyscale stimuli, created using 3 cat templates to 3 dog templates (Fig. 1A), two of which were randomly chosen for each participant. Stimuli were created by linearly combining one cat and one dog template, with changing contributions (morph level, e.g. 93.4% cat, 6.6% dog, with steps of 6.6). These stimuli have been used before on non-human primate research, and for more detail on their generation see ^43^. For each combination of cat and dog templates, 16 stimuli were created (8 dog + 8 cat stimuli). Each stimulus was categorized as either cat or dog depending on which category contributed more to the image (>50%). This yielded 32 cat and 32 dog stimuli, which differed in their distance to the category boundary (choice difficulty). We then added random Gaussian noise to these images, making classification more difficult (noise level). The amount of added noise was adapted to each participant using a staircase procedure (see below). This resulted in a 2 (categories) × 8 (choice difficulties) × 2 (noise levels) × 4 (template combinations) design with 128 unique stimuli. In order to increase the signal-to-noise ratio and have more trials in each cell of the design matrix, we then collapsed across all template combinations, and collapsed data from eight to four choice difficulty levels.

#### 4.2.5 Procedure

The experiment was programmed using PsychoPy3 (v.2020.1.3, ^44^). Participants started by performing a short training session outside the MR scanner, where they received trial-by-trial feedback on their responses. They first learned to classify template images, followed by classifying clean morphed images. They then entered the MR scanner, where they completed a staircase procedure to calibrate the noise level in the scanning environment. For this purpose, we presented template stimuli with varying levels of random Gaussian noise added. After each correct response, noise increased. After each wrong response, noise decreased. After seven reversals of noise change direction (up → down, down → up), the staircase procedure stopped and we applied the final noise level to all noisy stimuli used in the experiment.

After that, participants performed 6 runs of 128 trials of the experimental task. In each trial, a stimulus was presented on screen for 1.8 sec (Figure 1B), which was followed by a variable, pseudo-exponentially distributed inter-trial-interval (ITI, mean duration = 2.7 sec, range between 0.8 sec to 10.4 sec). Participants were instructed to respond while the stimulus was presented on screen, as quickly and accurately as possible. As an additional incentive, participants that were among the top 20% fastest *and* top 20% most accurate received a 10€ bonus payment. Participants responded using the left and right index fingers, using MR-compatible response boxes. Category-button-mappings were counter-balanced across runs for each participant (dog: right, cat: left in half of the runs, dog: left, cat: right in the other half). Participants received feedback at the end of each run (mean error rate + mean reaction time).

In each run, each unique stimulus was presented once, resulting in 8 repetitions of each combination of category (2), choice difficulty (4), and noise level (2). Category, choice difficulty, and template combination were pseudo-randomized and changed on a trial-by-trial basis. Noise level was blocked, and each run consisted of 4 blocks of 32 trials each. Two blocks were noisy, two blocks were clean, and half of the runs started with a noisy block, while the other half started with a clean block. Each block started with an instructions screen presented for 5 sec (‘Easy block starting, ’Hard block starting). We then ensured that none of the design variables was correlated, and that there were no sequential dependencies between trials, using a mutual information criterion (using permutation tests, p > 0.005).

Due to a coding error, for the first 29 participants, designs contained different numbers of cat and dog stimuli in each run, for each combination of choice difficulty and noise level. This introduced additional noise to the data, and made signal estimation in some conditions and some runs more difficult. However, due to the mutual information testing we employed before accepting designs, we ensured that this did not introduce any systematic biases even within participants.

### 2.3 Image acquisition

Functional imaging was performed on a 3T Siemens Prisma MRI scanner (Siemens Medical Systems, Erlangen, Germany), using a 64-channel head coil. For each of the six functional scanning runs, we acquired 350 T2*-weighted whole-brain echo-planar images (EPI, TR = 1730 ms, TE = 30 ms, image matrix = 84 × 84, FOV = 210 mm, flip angle=66°, slice thickness = 2.5 mm, voxel size = 2.5 × 2.5 × 2.5 mm, distance factor=0%, 50 slices with slice acceleration factor 2). Slices were oriented along the AC-PC line for each participant. A T1-weighted structural scan was acquired prior to the functional scans (MPRAGE, TR = 2250 ms, TE = 4.18 ms, TI = 900 ms, acquisition matrix = 256 × 256, FOV = 256 mm, flip angle=9°, voxel size = 1 × 1 × 1 mm). We further acquired 2 field maps (phase and magnitude) to correct for inhomogeneities in the magnetic field (TR = 520 ms, TE1 = 4.92 ms, TE2 = 7.38 ms, image matrix = 70 × 70, FOV = 210 mm, flip angle=60°, slice thickness = 3 mm, voxel size = 3 × 3 × 2.5 mm, distance factor=0%, 50 slices).

### 4.4 Analysis: Behavior

Behavioral data were analyzed using RStudio (version 1.2.1335, R version 4.0.3). We first removed all trials on which the participant failed to respond. On average, we removed 1.66% (SD = 0.64%) of all trials for each participant this way. Additionally, we removed trials with RTs < 150ms, removing 1.68% of trials on average (SD = 0.64%). To assess task performance, we extracted mean RTs and error rates for each combination of noise level and choice difficulty. For the RT analysis, we only used correct responses. RTs were then entered into a Bayesian ANOVA (*BayesFactor::anovaBF*, using the default scaled inverse chi-square prior on main effects and interactions, scaling factor fixed effects = 0.5, scaling factor random effects = 1), testing for evidence for or against both main effects and their interaction. Participants were entered as a random effect into this model. We interpreted the resulting Bayes Factors (BF10) according to ^45^. The same procedure was then applied to error rates.

We additionally fitted psychometric functions to the choice data, separately for each participant (Weibull function, using *quickpsy*, ^46^). Specifically, we computed the probability of choosing the dog response separately for each combination of morph level and noise level, and then fitted psychometric functions separately for both noise levels. This allowed us to extract several key parameters from the choice data: k, guess rate, and threshold. k determines the slope of the psychometric function and describes how sharply both categories are distinguished. The guess rate quantifies how often participants guess, and we expected k to be lower and the guess rate to be higher on noisy, as compared to clean, trials. To test this hypothesis, we entered estimates into a Bayesian paired t-test (*BayesFactor::ttestBF*, Cauchy prior, scaling factor = 0.707), comparing parameter values across noise levels. The threshold quantifies at which point on the scale of morphed images, ranging from 100% cat to 100% dog, participants were equally likely to choose cat or dog. We expected this to fall in the middle of the scale, i.e. where stimuli are close to 50% cat / 50% dog, and tested this hypothesis using a Bayesian t-test (Cauchy prior, scaling factor = 0.707).

### 4.5 Analysis: fMRI

fMRI analyses were performed using Matlab (R2018b, version 9.5.0.944444, The Mathworks), SPM12 (https://www.fil.ion.ucl.ac.uk/spm/), The Decoding Toolbox (v 3.99 ^47^), and RStudio (version 1.2.1335, R version 4.0.3). Raw data were first unwarped, realigned, and slice-time corrected (code: https://github.com/CCN-github/fMRI-preprocessing-SPM12). We then estimated normalization fields for each participant, which were then used to project mask files from normalized to native space. No spatial smoothing or normalization were applied to BOLD data to preserve fine-grained voxel activation patterns.

#### 4.5.1 First-level GLM analysis

Preprocessed data were used to estimate a voxel-wise general linear model (GLM ^48^). Sixteen regressors of interest were used, one for each combination of category (cat, dog), noise level (clean, noisy), and choice difficulty (1,2,3,4). We then added a variable number of nuisance regressors for each participant. First, we added condition-specific error regressors, modelling error trials separately for each condition. This led to a variable number of nuisance regressors, since not every run had errors in each condition. We chose condition-specific error regressors over a single error regressor, since we expected errors in very easy trials to derive from different processes than in very difficult trials (e.g. momentary lapse in attentional processes vs guessing). Second, we added six movement regressors. Regressors were time-locked to the onset of the stimulus presentation. We used the finite impulse response function as a basis function (FIR, 5 time bins with a duration of 1.73 sec each). This makes fewer assumptions about the shape of the haemodynamic response, compared to a canonical haemodynamic response function, making it better suited to model responses to short events in a heterogeneous set of brain regions from visual to prefrontal cortex (see ^49^ for a similar approach).

#### 4.5.2 Feature selection

##### ROI selection

Similar to ^32^, we defined a number of a-priori volumetric multiple demand (MD) regions-of-interest (ROIs), based on the Human Connectome Project atlas. The following regions were included in this experiment: SCEF, 8BM, 8C, IFJp, p9_46v, a9_46v, i68, AVI, IP1, IP2, PFm (see Figure 4, code: https://github.com/davidwisniewski/fmri-extract-HCP-mask). As an additional region of interest, we used the ventral visual cortex, as defined using the Harvard - Oxford atlas. Data from the chosen ROIs was extracted in native space for each subject, by projecting the ROI masks from MNI to native space separately for each participant, using the inverse normalization fields estimated during pre-processing.

##### Time-bin selection

Given that we use the FIR basis function, each regressor is modelled at five different time points. To select a time window of interest, we first estimated the haemodynamic lag to be 2TRs (3.46 sec), based on previous research using MVPA methods to extract task-related information from frontoparietal cortex ^49–51^. We then corrected for haemodynamic lag by using data from time bins 3 and 4, which started 3.46 sec and 5.19 sec after stimulus onset, respectively, for all multivariate pattern analyses. Again, this follows past research on time-resolved pattern analysis of task-related information ^52^. We expected haemodynamic responses to be short, given that the trial duration / stimulus processing was short, and this procedure strikes a balance between accounting for the expected short duration, and still allowing for the peak response to occur within a variable time window (between 3.46 sec and 6.92 sec after stimulus onset).

#### 4.5.3 Representational similarity analysis

For each ROI, we first extracted the beta weights for the 16 regressors of interest in each run. We then used The Decoding Toolbox ^47^ to perform a representational similarity analysis (RSA ^29,30^), using run-wise cross-validated Euclidean distance measures and applying multivariate noise normalization ^53^. Using cross-validated distances ensures that estimates are unbiased and average to zero if there is no systematic relation between activation patterns ^54^. All computed distances were then converted to a 16×16 representational distance matrix (RDM), representing pairwise distances between all conditions. This procedure was performed separately for each ROI, and for each of the two time bins of interest. For each ROI, we then averaged the RDMs across both time bins.

#### 4.5.4. Model-based RSA

The main goal of this experiment was to identify whether and how stimulus information is encoded in the MD network. For this purpose, we defined two alternative theoretical models of how stimulus information could be encoded in the brain.

First, stimulus information can be encoded in an equidistant format, i.e. all distances in the perceptual space can have an equivalent representation in neural representational space (Figure 3). Here, the full perceptual space is preserved in neural space, e.g. if a stimulus is 6.6% more ‘doggy’ than another stimulus, that distance will be half as large as to a stimulus that is 13.2% more ‘doggy’ in neural state space, irrespective of whether the two compared stimuli cross the decision boundary or not. This model assumes that slight perceptual differences between individual cat (and dog) stimuli are preserved, despite not being relevant for in the categorization task whatsoever. This model also contains category information, since the average between-category distance will be higher than the average within-category distance. Second, stimulus information can be encoded in a clustered, i.e. all distances between individual stimuli within a category (all dogs or all cats) are very small, while distances between categories are large. Here, coding of slight perceptual distances between individual stimuli within a category is dropped, and only the behaviorally relevant feature of the stimulus (category) is encoded.

In order to determine whether the models explain variance in the data, we first computed canonical, zero-order Spearman correlations between each model and the data, separately for each participant. For this purpose, we only considered data from the lower half of the RDM, excluding the diagonal. These correlations were Fisher-z transformed and entered into a Bayesian one-sided t-test (*BayesFactor::ttestBF*, Cauchy prior, scaling factor = 0.707) to assess whether they were above zero. This analysis was performed for each ROI separately, as well as separately for clean and noisy trials. Note that in principle, model-based RSA allows us to independently investigate evidence for both models, i.e. both, only one, or no model could potentially explain the data. Thus, even if both models partly explained the data, we would be able to identify this pattern of results.

Yet, in order to test our key hypotheses, this analysis was necessary but not sufficient. Zero-order correlations do not control for potential correlations between both explanatory variables / models, and the equidistant and clustered models were positively related here (r = 0.66). Thus, we repeated the same analysis described above, only now using partial Spearman correlations. This allowed us to quantify unique shared variance between the equidistant coding model and data RDMs extracted from visual cortex, while controlling for shared variance with the clustered coding model (and vice versa). This partial correlation analysis was again repeated separately for each participant, ROI, and noise level, and partial correlation coefficients were then Fisher-z transformed and entered into a Bayesian one-sided t-test (*BayesFactor::ttestBF*, Cauchy prior, scaling factor = 0.707) to assess whether they were above zero. To directly compare e.g. equidistant coding in clean and noisy trials, we computed paired Bayesian t-tests (*BayesFactor::ttestBF*, Cauchy prior, scaling factor = 0.707) between clean and noisy trials.

#### 4.5.5 Hypothesis 1: Visual cortex uses an equidistant coding format

To test this hypothesis, we performed a model-based partial correlation analysis, using the approach described above and data extracted from the ventral visual cortex. We expected the partial correlation between the equidistant model and the data to be above zero on clean trials (one-sided t-test vs 0), and further expected this correlation to be lower on noisy than on clean trials (one-sided paired sample t-test). We did not expect to see evidence for partial correlations between the clustered model and the data RDM.

#### 4.5.6 Hypothesis 2: Weak stimulus coding on clean trials in MD regions

On clean trials, we expected to see either no or weak equidistant stimulus coding in MD regions. This hypothesis was tested applying the same partial correlation approach as in Hypothesis 1 (one-sided t-tests vs 0) to each MD ROI separately. Again, we performed a control analysis explicitly testing for clustered coding as well.

#### 4.5.7 Hypothesis 3: Stronger, clustered coding on noisy trials in MD regions

Lastly, we hypothesized that stimulus coding would be stronger and more clustered on noisy trials in MD regions. To test this hypothesis, we again used our partial correlation approach to test whether clustered coding explains data on noisy trials in MD ROIs (one-sided t-test vs 0). In order to directly compare results across noise levels, we used paired one-sided t-tests. As a control analysis, we used the same approach to quantify evidence for equidistant coding on noisy trials as well.

#### 4.5.8 Exploratory analyses: Brain-behavior relations

In order to assess whether stimulus coding was related to performance and choice behavior, we performed a number of additional exploratory analyses. We first extracted the parameters of the psychometric function (slope, guess rate), separately for each participant. Then, we extracted the corresponding Fisher-z transformed partial correlations between the equidistant coding model and the data RDM. Both values were used to compute a Bayesian Pearson correlation (*BayesFactor::correlationBF*, Jeffreys prior, r = 0.33) across participants. This procedure was performed separately for clean and noisy trials. To assess differences between correlation coefficients, we computed the 95% credible interval (95% CI) for one correlation, and then tested whether the other correlation fell within this CI. If it did not, this was evidence for a difference between both coefficients.

## Acknowledgments

We would like to thank David Freedman for kindly sharing the stimulus material used in this study. DW was supported by the European Union’s Horizon 2020 research and innovation programme under the Marie Skłodowska-Curie grant agreement no. 665501 and the Flemish Science Foundation (FWO, FWO.KAN.2019.0023.01). CGG was supported by the Spanish Ministry of Science and Innovation (IJC2019-040208-I). SF was supported by the Einstein Foundation Berlin. AW was supported by Medical Research Council (U.K) intramural funding SUAG/052/G101400. MB was supported by an Einstein Strategic Professorship of the Einstein Foundation Berlin, and a GOA of the Special Research Fund of Ghent University (BOF.GOA.2017.0002.03).

## Author contributions

DW developed the study concept and analysis pipeline, performed the research, analyzed the data, interpreted the data, wrote the article draft, revised the draft, and funded the research. CGG developed the study concept and analysis pipeline, interpreted the data, and revised the draft. SF developed the study concept, interpreted the data, and revised the draft. AW developed the study concept, interpreted the data, and revised the draft. MB developed the study concept, interpreted the data, and revised the draft.

## Supplementary Material

### Supplementary Analysis 1: Psychometric function thresholds

Using the psychometric functions fitted to the choice data from each participant, we estimated each participant’s threshold (i.e. the point at which choosing cat and dog was equally likely) to determine whether choices were biased towards either category. Morph level ranged from 1 (0% dog) to 16 (100% dog), with the decision boundary being between 8 (most strongly morphed cat) and 9 (most strongly morphed dog). For unbiased choices, thresholds should thus be between 8 and 9 on the morph level scale. To test this hypothesis, we first computed a Bayesian t-test (*BayesFactor::ttestBF*, Jeffreys prior, r = 0.707) using data from clean images only.

We then extracted the 95% credible interval from the posterior distribution (number of iterations = 100.000), and tested whether this interval includes any values between 8 and 9. If it did, this would indicate that thresholds are indistinguishable from the expected threshold. If it did not, this would indicate that thresholds differ from the expected threshold. For clean trials, the 95% credible interval (95% CI = [8.21, 8.73]) indicated that thresholds were indeed between 8 and 9, as expected. For noisy trials, thresholds were somewhat lower and had a higher variance (95% CI = [7.10, 8.61]), but were still overlapping with the range of expected values. Using different priors (r = 1, r = 1.41) did not change this result. Thus, we concluded that choices were unbiased on clean trials, and that adding noise to the stimuli did not introduce biases.

### Supplementary Analysis 2: Simulations

Although the two model RDMs both capture stimulus information, they differ significantly in how stimulus categories are encoded. The clustered coding model assumes binary coding of stimulus information, discarding all differences between exemplars and only encoding category differences. The equidistant coding model however assumes that information about different exemplars is preserved in the neural code, and thus contains information about both categories and exemplars.

One effect of these assumptions is that the clustered coding model has an overall lower complexity, with the equidistant model making subtler predictions about representational distances even within the same category. Together with the fact that the equidistant coding model contains exemplar + category information while the clustered coding model contains only category information, this might lead to an unfair competition between these models in the partial correlation analyses used here. The clustered model might be ‘nested’ within the equidistant model, in that it does not contain unique information that is not also contained in the equidistant model. If this were the case, we would expect the clustered model to always be outperformed by the equidistant coding model, which might explain the lack of clustered coding we found.

To test this possibility, we performed a simulation analysis. Specifically, we generated data from either the clustered or equidistant model, and then tested whether our analysis approach was suitable to recover which model generated the data. To do so, we first took the clustered model RDM (range of values = [0,1]), and added varying amounts of random Gaussian noise to each cell of the matrix (mean = 0, sd = [0.01 - 3.00]). This procedure was repeated 1000 times to generate 1000 simulated ‘participants’. These simulated RDMs were then used as input to the same analyses performed on the actual data in the main manuscript, including both the zero-order/canonical and partial correlation analyses. We expected correlations of the simulated RDMs and the clustered model to be higher than with the equidistant coding model, which would indicate successful model ‘recovery’.

Results (Figure S2 A.) indicated that the clustered model better explained the data regardless of noise level in the zero-order correlation analysis, although the equidistant model also explained simulated data to some degree. In the partial correlation analysis, it can be clearly seen that only the clustered model explained the data, while the equidistant model was effectively suppressed and unrelated to the data, irrespective of how noisy the data was assumed to be. We repeated the same analysis, only generating data from the equidistant coding model, and results showed the equidistant model explaining the simulated data better than the clustered model (Figure S2 B). Thus, using our analysis approach, we could recover which model was used to generate the simulated data well, regardless of how noisy data RDMs are assumed to be.

One might argue that these analyses incorrectly assumed that data RDMs are either 100% clustered or 100% equidistant. In reality, data RDMs likely reflect a mix of clustered and equidistant signals, which we did not account for in the above analyses. But what if both models contributed equally to the signal? In this case, both models should explain the data equally well, and this seems a more likely scenario than assuming data are either 100% clustered or 100% equidistant. To test this, we first generated data similarly to the previous two simulations, only now using a ‘mixed’ model in which both the equidistant and clustered models contribute equally. This mixed model was computed by taking the mean of both model matrices, i.e. in each cell of the matrix both models contributed to an equal degree. In addition to being more realistic, this analysis was also more sensitive to detect potential biases towards either model in our analysis approach, as in principle both models should be able to explain an equal part of unique variance in the data in the partial correlation analysis. If one model outperformed the other, our analysis would be biased towards detecting it in our data. Simulation results (Figure S2 C) suggest that the clustered model explains the data slightly better when noise is weak. For noisier data, both models explain the data equally well. Thus, if anything, our analysis approach is slightly biased towards detecting clustered coding (assuming little noise), making our findings of equidistant coding in parietal cortex and other MD regions even more striking.

**Figure S1:**
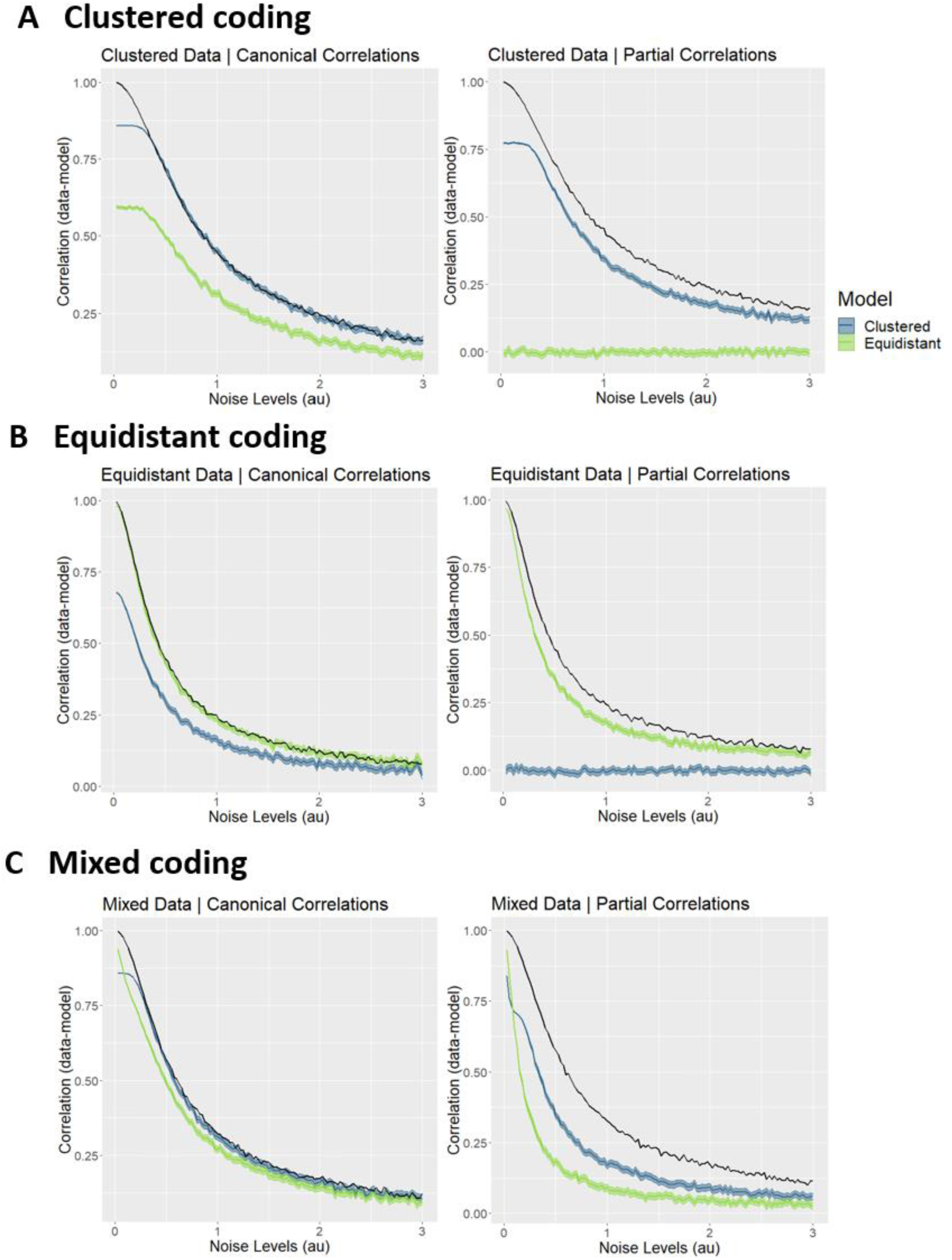
Simulation results. **A**. **Clustered data.** Correlations between simulated data, generated from the clustered coding model, and the two different model RDMs (clustered, equidistant), for different amounts of noise on the data RDMs. **B. Equidistant data**. The same correlations are depicted, only for data generated from the equidistant model. **C. Mixed data**. The same correlations are depicted, only for data generated from a mixed model (50% equidistant coding, 50% clustered coding). Zero-order correlations are depicted on the left, partial correlations on the right. Shaded areas represent 90% confidence intervals. The black line represents the noise ceiling, estimated following ^30^, and indicates the maximum correlation that can be expected given a particular level of noise in the data.

